# Landscape configurations determining the genetic structure of the Yellow-Spotted Amazon River Turtle (*Podocnemis unifilis*) in Brazilian Amazonia

**DOI:** 10.1101/2022.08.23.504961

**Authors:** Maria Augusta Paes Agostini, Arielli Fabrício Machado, Camila Duarte Ritter, Maria das Neves da Silva Viana, Luiz Alberto dos Santos Monjeló, Paulo César Machado Andrade, Jackson Pantoja-Lima, Juarez C. B. Pezzuti, Daniely Félix-Silva, Waldesse Piragé de Oliveira, Richard C. Vogt, Tomas Hrbek, Izeni Pires Farias

**Affiliations:** Programa de Pós-Graduação em Biodiversidade e Biotecnologia da Amazônia Legal – BIONORTE, Av. Carvalho Leal, 177, Manaus, AM, 69065-001, Brazil; Laboratório de Evolução e Genética Animal (LEGAL), Departamento de Genética, Universidade Federal do Amazonas (UFAM), Av. General Rodrigo Octávio Jordão Ramos, 1200, Coroado I, Manaus, AM, 69067-005, Brazil; Laboratório de Biotecnologia, Universidade Federal do Tocantins (UFT), Palmas, TO, 77001-090, Brazil; Centro de Estudos de Quelônios da Amazônia (CEQUA), Instituto Nacional de Pesquisa da Amazônia (INPA), Avenida André Araújo, 2936, Petrópolis, Manaus - AM, 69067-375, Brazil; Laboratório de Paleobiologia, Universidade Federal do Pampa (UNIPAMPA), BR 290 - km 423, Universitário, São Gabriel, RS, 97307-020, Brazil; ATGC Genética Ambiental LTDA. Rua dos Funcionários 1540, Juvevê, Curitiba, PR, 80035-050, Brazil; Grupo Integrado de Aquicultura e Estudos Ambientais, Departamento de Zootecnia, Universidade Federal do Paraná (UFPR), Rua dos Funcionários, 1540, Juvevê, Curitiba, PR, 80035-050, Brazil; Departamento de Fisiologia, Universidade Federal do Amazonas (UFAM), Av. General Rodrigo Octávio Jordão Ramos, 1200 - Coroado I, Manaus, AM, 69067-005, Brazil; Faculdade de Ciências Agrarias, Universidade Federal do Amazonas (UFAM), Manaus, AM, Brazil; Instituto Federal de Educação, Ciência e Tecnologia do Amazonas (IFAM), Presidente Figueiredo, AM, Brazil; Núcleo de Altos Estudos Amazônicos (NAEA), Universidade Federal do Pará (UFPA), Belém, PA, 66075-110, Brazil

**Keywords:** Freshwater turtle, Landscape Genetics, mtDNA, Population Structure, Conservation

## Abstract

**Context:** Waterfalls and rapids of Amazon basin have been suggested as causing the speciation and genetic structure of many freshwater species, including turtles. The species behavior affects the way waterfalls and rapids limit gene flow. The Yellow-spotted River Turtle (*Podocnemis unifilis*), a widely distributed and endangered Amazonian turtle, does not show the habit to migrate long distances for breeding or eating, but has a complex geographic pattern of genetic variation.

**Objectives:** Here, we investigate isolation by distance and by resistance of *P. unifilis*. We analyzed if the species ecological niche and waterfalls explain the genetic distance in Brazilian Amazonia.

**Methods:** We evaluated the *P. unifilis* spatial distribution of genetic variability and diversity using the control region of mitochondrial DNA. We tested the hypotheses of isolation either by distance and resistance through an integrative approach using genetic, geographic, and ecological data. We created a resistance matrix using species niche modeling. We compared the explanation power of geographical distance (both linear and in-water distance) and resistance distance on genetic distance (ΦST fixation index) using multiple regressions and Mantel tests.

**Results:** We found a high genetic diversity and pattern of genetic structure proved to be geographically complex. The population structure followed some watersheds but also showed structuring within different rivers. We found that landscape resistance better explains genetic distance than linear and in-water distance.

**Conclusions:** The resistance of the landscape influences the displacement of individuals by aquatic, vegetational, biological, and geomorphological variables, and efforts to species conservation need to be applied throughout its distribution considering landscape genetics.

## Introduction

The Amazon basin is the world’s biggest basin and their water and landscape structure affect the gene flow of species in the region (Hoorn et al. 2010). For instance, the abrupt transition from the crystalline Guyana and Central Brazil shields to the sedimentary plain marks the landscape in the form of waterfalls and stretches of rapids in the largest tributaries of the Solimões-Amazon rivers (Ayres 1995). Those geomorphological structures can play a role of natural hurdle to gene flow and/or as agents of diversification. In some places, rapids and waterfalls are seen as important elements in the historical processes of allopatric speciation, population structure, and even limiting gene flow in some aquatic and semiaquatic groups (Willis et al. 2012a; Muniz et al. 2018), varying with the migratory capacity of the species.

In the Madeira river was possible to identify the delimitation of fish species of the genus *Cichla* (Willis et al. 2012a). Also, its series of 18 rapids limits the gene flow unidirectionally of dolphin (*Inia geoffrensis*) populations (Gravena et al. 2014), and restricts, but does not prevent the migration of the largest South America freshwater turtle (*Podocnemis expansa*) (Pearse et al. 2006). Elsewhere, waterfalls and rapids have been suggested as causing the population genetic structure of a low-dispersion turtle (*Podocnemis erythrocephala*) (Santos et al. 2016).

As Amazonian turtles have different migration capability and feed behavior (Ferrara et al. 2017), they are an ideal group to study the effects of landscape in their genetic structure (Oliveira et al. 2019). *Podocnemis expansa* is adapted for swimming hundreds of kilometers, but genetic differences were found only between sub-basins (Pearse et al. 2006). In contrast, the Yellow-spotted River Turtle (*Podocnemis unifilis*) that is widely distributed (Ferrara et al. 2017; Turtle Taxonomy Working Group 2021), does not show the habit to migrate long distances for feeding or nesting (Vogt 2008; Andrade 2012, 2015; Ferrara et al. 2017), although this species can walk on land considerable distances (Almeida et al. 2006, Alcântara et al. 2013), possibly have a geographic pattern of genetic variation associated with landscape features (Escalona et al. 2009).

Despite *P. unifilis* was considered an Endangered species in the professional Red List of the Tortoise and Freshwater Turtle Specialist Group – TFTSG (Turtle Taxonomy Working Group 2021) and Vulnerable in the International Union for Conservation of Nature and Natural Resources (IUCN, 2016) Red List, a serious paucity of information on its genetics persists. Indeed, in some Western Amazonia drainages, the gene flow occurs only through river channels, while in others, individuals can migrate across wetlands and small watercourses (Escalona et al. 2009). However, gene flow patterns between populations of this species in the rest of the Amazon Basin remain unknown.

Although dispersion allows gene flow and reduces inbreeding (Ronce 2007), it has a price. Dispersions in a non-suitable matrix have high energetic costs and mortality risks by biotic, abiotic, and/or anthropogenic features (Fahrig 1998; Gruber and Henle 2008; Ruiz-González et al. 2015). Thus, it is possible that geomorphological structures like waterfalls and rapids, or even other ecological and landscape characteristics, limit gene flow of *P. unifilis* throughout most of its distribution, as has been seen in other turtle species in the Amazon basin (Sites et al. 1999; Pearse et al. 2006; Vargas-Ramírez et al. 2008; Santos et al. 2016; Michels and Vargas-Ramírez 2018).

To test which environmental variables can influence *P. unifilis* migration, it is important to consider different factors to link the landscape arrangement with the genetic variation of the species (DiLeo and Wagner 2016). When no or little landscape configuration determines gene flow, it is expected an “isolation by distance” (IBD, Wright 1943; Van Strien et al. 2015). In contrast, when physical natural barriers or non-suitable matrix types difficult the gene flow, “isolation by resistance” (IBR, McRae 2006) is expected. Therefore, it is necessary to verify the influence on population genetic patterns by the ecological characteristics of the landscape, as well by climate, geology, and geomorphology, focusing on river systems (Oliveira et al. 2019).

Here, we used the molecular marker of the mitochondrial DNA (mtDNA) control region to assess the spatial distribution of the genetic variability and diversity of *P. unifilis* along the main rivers of the Brazilian Amazonia. We characterized population structure and tested isolation by distance (IBD) and by resistance (IBR) to verify if landscape features can explain the genetic distance of *P. unifilis*.

## Material and Methods

### Sample collection, DNA extraction and molecular data

We sampled a total of 370 adult turtles from 22 localities at different rivers (some of them upstream and downstream of waterfalls and rapids), in the Brazilian Amazon basin (Fig. 1). The sequences of the *P. unifilis* database from the Animal Evolution and Genetics, and Molecular Biochemistry Laboratories, both at the Federal University of Amazonas (UFAM) in Manaus, State of Amazonas, Brazil, are from samples collected between 2001 to 2018 by different groups of field researchers.

**Fig. 1.**
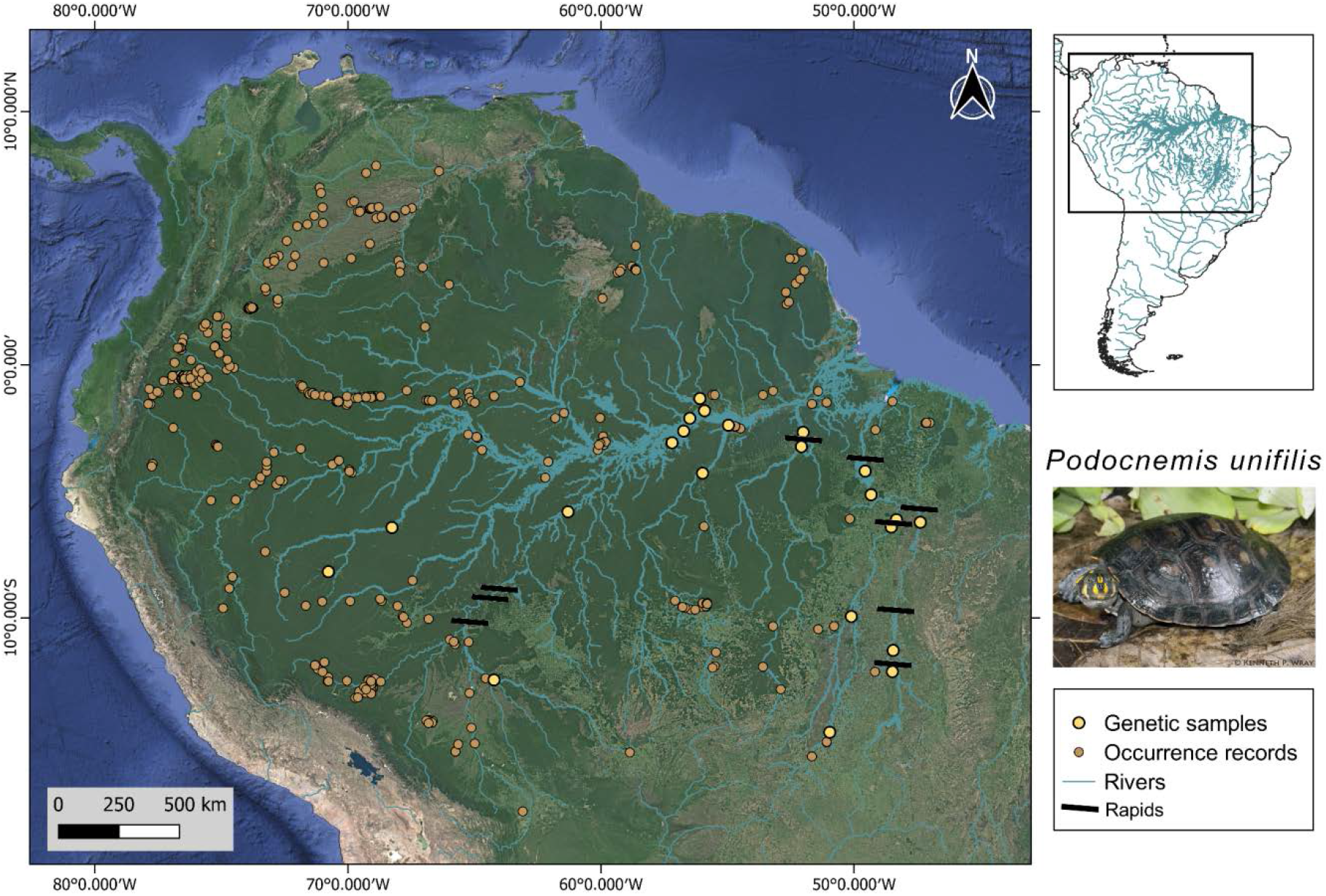
Map of the distribution *Podocnemis unifilis*, the Yellow-Spotted River Turtle. Points in light yellow with thicker edges are genetic samples sites, points in dark yellow with thinner edges are the occurrence records, blue lines are the rivers and black bars are the rapids. Photo by Kenneth P. Wray.

Turtles were captured with fyke nets, trammel nets, fishing or hand capture on nesting beaches depending on the region (Vogt 2016). Geographical coordinates were taken at capture locations with a Garmin GPS meter. All turtles captured were marked with a unique code notching the marginal scutes (Cagle 1939). Blood samples were taken by tail or femoral vein puncture and stored in 96% alcohol in a cooler or refrigerator at 4° C until the processing.

Total genomic DNA was extracted using CTAB method and modified protocol (Doyle and Doyle 1990). The amount of DNA per sample was measured in the NanoDrop 2000 spectrophotometer (ThermoScientific). We amplify the mtDNA control region with the primer pair: PRO (5’-CCCATCACCCACTCCCAAAGC-3’) (Pearse et al. 2006) and 12SR5 (5’-GTCAGGACCATGCCTTTGTG-3’) (Kocher et al. 1989). Each tube contained 4 mm MgCl2, 0.8 mM dNTPs, 1.0x enzyme Taq polymerase buffer (100 mM Tris-HCl, pH 8.5, 500 mM KCl), 1pMol of each primer, 1.5 U Taq DNA polymerase enzyme and 50ng DNA supplemented with water to a final volume of 15 μL. PCR reactions were submitted to the following cycling profile: hot start at 94° C for 1 minute, followed by 35 cycles of denaturing at 94° C for 1 minute, annealing at 58° C for 1 minute and extension at 72° C for 1 minute and 30 seconds, and final extension was performed at 72° C for 5 minutes.

Amplified samples were purified using Thermo Scientific Exonuclease enzymes I and Thermo Scientific Fast AP Thermosensitive Alkaline Phosphatase (ThermoScientific) for 30 minutes at 37° C followed by 15 minutes at 80° C. The amplifications were subjected to sequencing reaction using the Big Dye Terminator Cycle Sequencing Kit (ThermoScientific), using the internal primer DLSex (primer 5’-AGTGCTCTTCCCCATATTATG-3’) (Viana et al. 2017) according to the manufacturer’s recommendation and resolved on an ABI 3500 automated sequencer (Thermo Fisher Scientific, Brazil).

The sequences were edited and aligned in the Geneious version 6.0 software (Kearse et al. 2012) using the ClustalW tool (Thompson et al. 1994) under default conditions and reviewed manually.

### Genetic diversity, neutrality tests and distribution of haplotypes

For each locality we estimated the numbers of haplotypes (nH), segregating sites (S), haplotype (Ĥ), nucleotide diversity (Π) (Tajima 1983; Nei 1987), *D* (Tajima 1989), and *Fs* (Fu 1997) neutrality tests, in the program Arlequin version 3.5 (Excoffier and Lischer 2010).

To visualize the haplotype network, we generated a maximum likelihood tree using the evolution model HKY [{3, 1, 1, 1, 1, 3}, Empirical]) inferred in the program TreeFinder v. 2011 (Jobb et al. 2004). The maximum likelihood tree was then converted in the genealogies of haplotypes through the HaploViewer program (Salzburger et al. 2011).

### Population structure and gene flow

The proportions of genetic differentiation between the sampled areas were calculated using the analysis of molecular variance – AMOVA (Excoffier et al. 1992) with the significance levels estimated using 10,000 permutations, implemented in the program Arlequin. We used the Bonferroni correction (Rice 1989) for multiples comparisons. Fixation indexes ΦST were used to estimate indirectly the actual number of female migrants (equivalent to gene flow) using the equation Nefm = ((1/FST) - 1) (Weir and Cockerham 1984).

Population structure was assessed with the Bayesian Analysis of Population Structure in BAPS version 5.2 (Corander et al. 2008) and then we estimate the genetic diversity indices for each biological group in the program Arlequin version 3.5 (Excoffier and Lischer 2010).

### Ecological niche modeling for resistance landscape analysis

We generated an Ecological Niche Model (ENM) for *Podocnemis unifilis* to be used as a baseline landscape resistance model representing the gradients of environmental suitability for the species occurrence. We used digital layers of a continuous freshwater-specific set of environmental bioclimatic and soil variables extracted from the EarthEnv database (Domisch et al. 2015), aquatic variables from the HYDRO1k (USGS 2018) and a vegetation variable (Leaf Area Index, Myneni et al. 2015), all containing about 5 km of resolution. These variables were cropped considering the approximate limits of the species distribution according to the occurrence records (26°S 82°W, 14°N 36°W) using the raster package v. 3.4-13 (Hijmans and Van Etten 2021) of R v.4.1.0 software (R Core Team 2021). Species occurrence records were compiled from SpeciesLink (https://specieslink.net/), Global Biodiversity Information Facility (GBIF, https://www.gbif.org/) and Vertnet (http://vertnet.org/) databases, also including records from the Instituto Nacional de Pesquisas da Amazônia collection (Manaus, Amazonas, Brazil). These records were cross-checked with records considered in literature reviews by Amazonian turtle specialists (Ferrara et al. 2017; Rhodin et al. 2018), resulting in 3,415 occurrence records from 340 localities (Fig. 1, Table S1).

To reduce the effect of sampling bias on the model, we used a sampling bias file generated through a grid of probabilities based on species occurrence records, where the values of the sites reflect the variation in the sampling effort to weight the ENMs (Elith et al. 2011). The sampling bias file was generated using the kernel density estimation method with the ‘kernelUD’ function from the adehabitatHR v.0.4.16 package (Calenge 2006) in R software (R Core Team 2021). The digital layers of the cropped environmental variables were used to calibrate and test the model.

We evaluated the level of correlation between the variables by Pearson’s correlation coefficient test using the ‘cor’ function of the stats 3.6.2 package of R (R Core Team 2021) and following the same procedure used by Rissler and Apodaca (2007) which considers the correlation threshold of 0.75. Only the variables with the greatest biological relevance for the species were used to generate the final model by evaluating the results of the Jackknife graphs generated in the preliminary model where each variable is tested separately and excluding each other variable, revealing the gain and loss of models containing or not each variable (Phillips 2006).

The ENM of *P. unifilis* (Fig. S1) was built with all the non-duplicated occurrence records of the species (340 records, Table S1), the sampling bias file and previously selected uncorrelated environmental variables in MaxEnt v.3.4.4 (Steven et al. 2017). The ENM was generated using 10 independent replicates, default MaxEnt parameters (except for the betamultiplayer which was selected to 5.0 considering the wide species distribution), and the method of cross-validation of the points to evaluate the model. We chose the logistic output for the presentation of the model in the geographical space (potential distribution), with each pixel representing environmental suitability ranging from 0 (representing minimum environmental suitability) to 1 (maximum environmental suitability) (Phillips 2006). The performance of the model was assessed using the Area Under the Curve (AUC) method, considering the limit value of AUC > 0.7 to accept the model (Phillips 2006).

### Isolation by distance and by resistance

To test the isolation by distance, we built a geographic distance matrix, in which we calculated the pairwise minimal distances between all locations with the geosphere v.1.5.10 R package (Hijmans et al. 2019). Additionally, we calculated the pairwise in-water distance, that is the distance of one point to the other just considering the rivers paths calculated with the ruler tool from Google Earth (http://earth.google.com/).

To test the isolation by resistance, we created a landscape friction layer to calculate least environmental cost distance matrix by inverting the values of the environmental suitability based on ENM (which ranges from 0 to 1) using the raster v.1.5-10 R package (Hijmans and Van Etten 2021). Then, areas with greater environmental suitability for species occurrence (pixels = 1 in the ENM) present low resistance to displacement (pixels = 0.01 in the landscape friction layer) and areas with lower environmental suitability (pixels = 0 in the ENM) present high resistance to displacement (pixels = 0.99, in the landscape friction layer). We tested this model of resistance and then adjusted the values of pixels in the friction layer of the landscape including rapids and waterfalls located between localities with genetic sampling, the resistance values is proportional to its extension varying between 0.50 and 0.85. We tested this new model and started to use it for further analyses. (Table S2).

To generate the least environmental cost distance matrix we created a transition object with the gdistance R package v.1.1-4 (Van Etten 2012) using the friction layer used for the least cost paths (LCPs), an estimation of the pixel-to-pixel distances, and Moore’s neighborhood (directions = 16), consisting of all pixels surrounding the target pixel (Van Etten 2012). We then corrected the transition object, and finally, we calculated the least environmental cost distance between points, using the gdistance. We also generated a map showing the LCPs using the leastcostpath v.1.8.0 R package (Lewis 2021).

To determine if genetic distance is better predicted by minimum geographic distance, in-water distance, or resistance, we used the pairwise genetic distance (ΦST values) between localities as predictor and then we performed three Mantel tests for each of these explanatory variables (pairwise minimum geographic distance, pairwise in-water distance, and pairwise least cost distance) with the ade4 v.1.0.12 R package (Dray and Dufour 2007), with a permutation of 10,000 for significance.

Furthermore, we performed multiple regressions on dissimilarity matrices (MRM), as implemented in the function ‘MRM’ of the R package ecodist v.2.0.1 (Goslee and Urban 2007), using as response variable the genetic distances (ΦST) and as explanatory variables the minimum geographic distance, in-water distance, or the least cost distance.

## Results

### Genetic diversity and neutrality tests

We analyzed 430 pb of 370 individuals of *P. unifilis*, totaling 216 polymorphic sites, 138 haplotypes, and different levels of genetic diversity, with haplotype diversity (Ĥ) ranging from minimum in Pium (0.000) to maximum in Peixe (1.00) and nucleotide diversity (Π) ranging from minimum again in Pium (0.000) to the maximum nucleotide diversity in Itaituba (0.018) (Table 1).

**Table 1.** Localities sampled and genetic diversity indices for *Podocnemis unifilis*.

| # | Locality | Water body | County | Code | Latitude | Longitude | N | S | h | Hd | $\Pi$ | D | Fs |
| --- | --- | --- | --- | --- | --- | --- | --- | --- | --- | --- | --- | --- | --- |
| 1 | Tarauacá | Juruá | AC | JuruTara | 8°10'37.89"S | 70°46'18.06"W | 12 | 3 | 3 | 0.439 +/- 0.158 | 0.002 +/- 0.002 | -0.829 | 0.10495 |
| 2 | Itamarati | Juruá | AM | JuruItam | 6°25'55.27"S | 68°15'35.91"W | 12 | 3 | 3 | 0.318 +/- 0.164 | 0.001 +/- 0.001 | **-1.629** | -0.61396 |
| 3 | Manicoré | Madeira | AM | MadeMani | 5°48'37.17"S | 61°18'15.52"W | 16 | 12 | 7 | 0.775 +/- 0.088 | 0.009 +/- 0.005 | 0.106 | 0.08084 |
| 4 | Costa Marques | Guaporé | RO | MadeCost | 12°27'11.31"S | 64°13'42.04"W | 19 | 6 | 3 | 0.433 +/- 0.117 | 0.005 +/- 0.003 | 0.953 | 3.50430 |
| 5 | Parintins | Amazonas | AM | AmazPari | 2°37'13.67"S | 56°43'56.65"W | 22 | 9 | 6 | 0.736 +/- 0.070 | 0.008 +/- 0.005 | 1.262 | 1.51946 |
| 6 | Barreirinha | Amazonas | AM | AmazBarr | 3° 4'58.02"S | 57°11'29.39"W | 17 | 8 | 4 | 0.500 +/- 0.136 | 0.006 +/- 0.004 | 0.420 | 2.37282 |
| 7 | Terra Santa | Nhamundá | PA | NhamTerr | 2° 6'59.79"S | 56°29'28.82"W | 21 | 12 | 10 | 0.814 +/- 0.081 | 0.010 +/- 0.006 | 0.818 | -1.37269 |
| 8 | Oriximiná | Trombetas | AM | TromSapu | 1°49'1.47"S | 55°54'33.93"W | 10 | 15 | 8 | 0.933 +/- 0.077 | 0.014 +/- 0.008 | 0.567 | -1.52677 |
| 9 | Oriximiná | Trombetas | AM | TromJara | 1°20'6.60"S | 56° 5'25.80"W | 15 | 20 | 12 | 0.971 +/- 0.033 | 0.010 +/- 0.006 | -1.300 | -5.68576 |
| 10 | Santarém | Tapajós | PA | TapaSant | 2°23'23.21"S | 54°57'52.61"W | 8 | 13 | 6 | 0.929 +/- 0.084 | 0.013 +/- 0.008 | 0.527 | -0.27339 |
| 11 | Itaituba | Tapajós | PA | TapaItai | 4°17'16.52"S | 55°59'8.04"W | 9 | 15 | 4 | 0.694 +/- 0.147 | 0.018 +/- 0.011 | 1.975 | 4.00065 |
| 12 | Altamira | Xingu | PA | XingAlta | 2°40'8.21"S | 52° 0'47.66"W | 27 | 18 | 7 | 0.815 +/- 0.043 | 0.011 +/- 0.006 | 0.0404 | 2.34479 |
| 13 | Vitória do Xingu | Xingu | PA | XingVito | 3°14'11.02"S | 52° 4'53.50"W | 21 | 9 | 8 | 0.791 +/- 0.066 | 0.005 +/- 0.003 | -0.693 | -2.08899 |
| 14 | Tucuruí | Tocantins | PA | TocaTucu | 4°12'7.87"S | 49°33'43.48"W | 22 | 9 | 6 | 0.771 +/- 0.052 | 0.003 +/- 0.002 | -1.330 | -0.93697 |
| 15 | Itupiranga | Tocantins | PA | TocaItup | 5° 8'7.74"S | 49°19'40.14"W | 6 | 6 | 3 | 0.600 +/- 0.215 | 0.005 +/- 0.004 | **-1.367** | 1.01973 |
| 16 | São João | Araguaia | PA | AragAnto | 6° 7'5.96"S | 48°19'38.51"W | 28 | 17 | 10 | 0.762 +/- 0.074 | 0.005 +/- 0.003 | **-1.934** | **-3.52272** |
| 17 | Xambioa | Araguaia | TO | AragXamb | 6°24'52.54"S | 48°31'56.48"W | 35 | 5 | 5 | 0.393 +/- 0.096 | 0.001 +/- 0.001 | **-1.499** | **-2.22205** |
| 18 | Pium | Araguaia | TO | AragPium | 9°56'54.64"S | 50° 6'33.62"W | 6 | 0 | 1 | 0.000 +/- 0.000 | 0.000 +/- 0.000 | 0.000 | NA |
| 19 | Cocalinho | Araguaia | MT | AragCoca | 14°31'14.82"S | 50°57'44.57"W | 19 | 8 | 7 | 0.749 +/- 0.088 | 0.003 +/- 0.002 | -1.183 | **-2.23221** |
| 20 | Tocantinópolis | Tocantins | TO | TocaToca | 6°13'23.03"S | 47°22'58.69"W | 20 | 12 | 14 | 0.953 +/- 0.033 | 0.007 +/- 0.004 | -0.339 | **-8.34057** |
| 21 | Ipueiras | Tocantins | TO | TocaIpue | 11°17'15.19"S | 48°27'19.86"W | 20 | 5 | 6 | 0.632 +/- 0.113 | 0.002 +/- 0.002 | -0.774 | **-2.10491** |
| 22 | Peixe | Tocantins | TO | TocaPeix | 12° 7'35.62"S | 48°29'18.92"W | 5 | 11 | 5 | 1.000 +/- 0.127 | 0.011 +/- 0.008 | -0.654 | -1.41093 |
| *Sum* |  |  |  |  |  |  | **370** | **216** | **138** |  |  |  |  |
| *Mean* |  |  |  |  |  |  | 17 | 10 | 6 |  |  |  |  |

**Notes:** Localities numbers and codes are according to a map of samples. N: number of individuals per location. S: number of polymorphic sites. h: number of haplotypes. Hd: haplotype diversity. $\pi$ : nucleotide diversity; D= Tajima's D test and Fs= Fu's Fs test with p values ( $\alpha \leq 0.05$ ) in bold. County abbreviations: AM – Amazonas; AC – Acre; PA – Pará; RO – Rondônia; TO - Tocantins. Coordinates are in decimal degrees, datum WGS 84.

The neutrality tests were significant only in the localities of the Tocantins-Araguaia basin. The Tajima’s D test shows significant values in four localities (2: JutuItam, 15: TocaItup, 16: AragAnto and 17: AragXamb), and Fu’s Fs only in five (16: AragAnto, 17: AragXamb, 19: AragCoca, 20: TocaToca and 21: TocaIpue; Table 1), indicating that the remaining populations are evolving according to the hypothesis of selective neutrality in relation to haplotypes of the mtDNA control region.

### Population structure and gene flow

Bayesian analysis of population structure (BAPS) indicated the occurrence of four biological clusters (log [marginal likelihood] of optimal partition = - 2647.2048, 1.00 probability of K = 4) between individuals (Fig. 2).

**Fig. 2.**
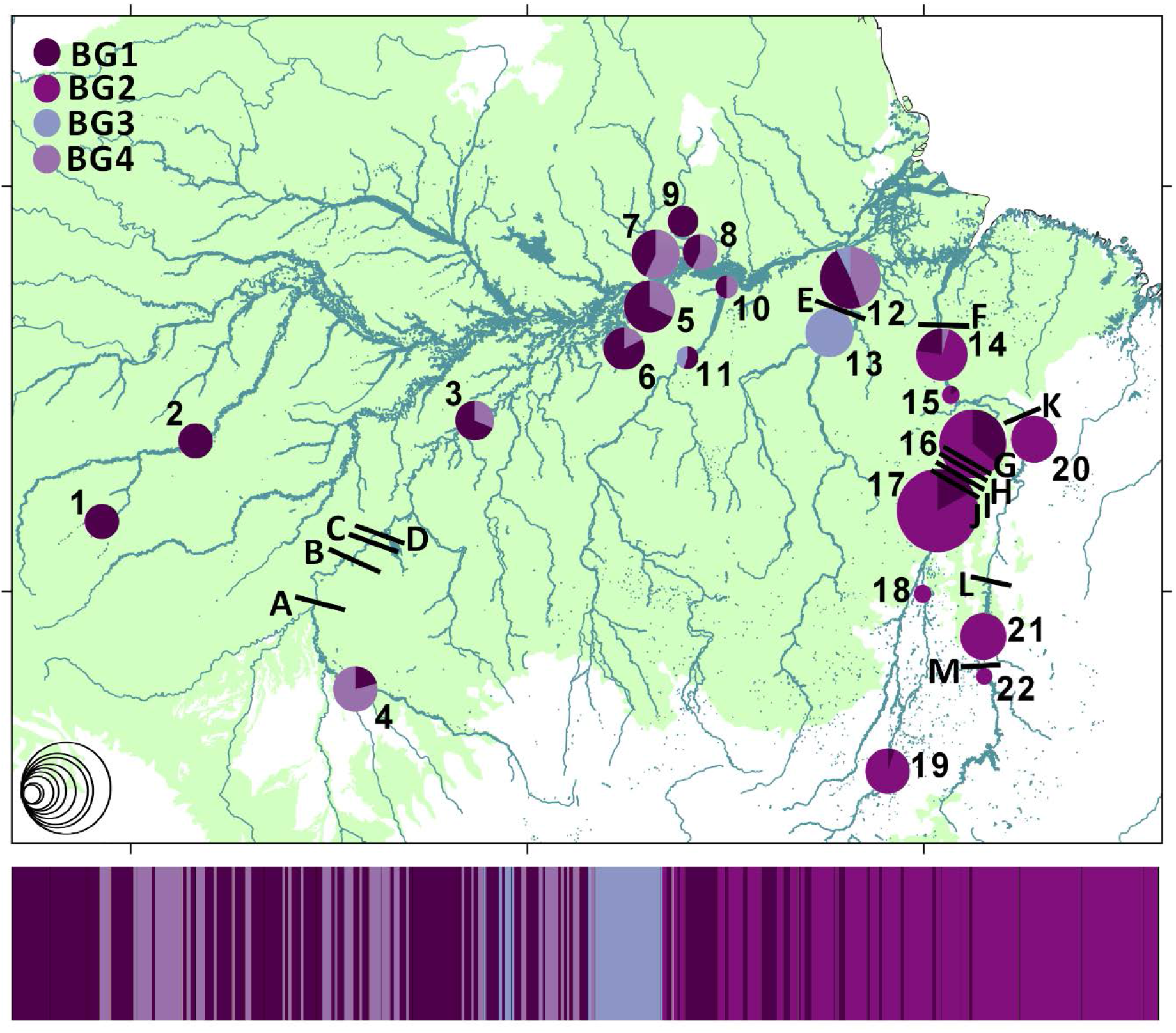
Map showing the geographic distribution of the biological groups of *Podocnemis unifilis* found with the Bayesian Analysis of Population Structure (BAPS). Letters represent waterfalls: A) Misericórdia, B) Caldeirão, C) Teotônio, D) Santo Antônio, E) Volta Grande do Xingu, F) Taboca, G) Santa Isabel, H) São João, I) Remanso dos Botos, J) São Miguel, K) Santa Rita, L) Lageado, and M) Arquipélago do Tropeço. **Notes:** The colors at each circle on the map represent the cluster to which the analyzed individuals belong based on BAPS assignments, the circle size represents the number of individuals sampled (ranging from 5 to 35 according to the scale at the bottom left), blue lines represent the rivers and black bars represent rapids and waterfalls.

The Biological Group 1 (BG1) has 152 individuals and was found in almost all sampled locations in the Brazilian Amazonia, distributed from the Juruá river to the Tocantins-Araguaia basin. BG2 is restricted to localities in the Tocantins-Araguaia river basin (126 individuals), not having been observed in any population of the Amazon river basin. The localite upstream of the Xingu (12: XingAlta) appears as a group (21 of the 27 individuals belong to the BG3; Fig. S2), which also has individuals in localities upstream of the Tapajós (11: TapaItai) and the Downstream of the Xingu (13: XingVito). BG4 emerged in the Madeira river basin and is distributed throughout the central region of the Amazon river basin to the downstream streams and waterfalls of the Trombetas, Tapajós, Xingu and lower Tocantins rivers (Fig. 2; S2). Analyzing the genetic diversity of the biological groups (Table S3) we found high values of haplotype diversity (Ĥ) ranging from 0.776 (BG4) to 0.886 (BG3) and nucleotide diversity (Π) ranging from 0.005 (BG2 and BG4) to 0.006 (BG1 and BG3).

Of the 138 haplotypes sampled, 75 were unique and four were more frequent with individuals from several localities (Fig. S2). The greatest sharing of haplotypes was between 61 individuals from the Juruá, Madeira, Amazonas, Tocantins and Araguaia rivers, followed by 51 and 29 samples only from the Tocantins-Araguaia basin. We highlight here one haplotype with the most of the 14 samples from municipality of Barreirinha in the State of Amazonas, and another with 14 individuals only from Guaporé river (Fig S2).

Analysis of molecular variance (AMOVA) showed that 50.95% of variation is distributed among localities and 49.05% within each locality, indicating a high degree of genetic population subdivision between the locations sampled (Φst = 0.51, p < 0.0001). It is also checked when analyzed the pairwise comparison of Φst, where about 94% of the comparisons were significant, and 77% after Bonferroni correction (lower diagonal of the Table S4).

Higher and significant Φst values were observed for comparisons between localities 4: MadeCost, 9: TromJara, 11: TapaItai, 13: XingVito, 14: TocaTucu, 20: TocaToca and 22: TocaPeix, and 6: AmazBarr (Table S4). This genetic structure can also be seen in other analyzes (Bayesian; Fig. 2, and haplotype genealogy; Fig. S2), involving the locations above the waterfalls and rapids, specially at in the Guaporé, Xingu, and Tocantins rivers.

Analyzing the estimated number of female migrants (Nefm) per generation between localities studied, we found that most of localities are structured with values < 1 (55%). Twenty four percent of the comparisons showed restricted gene flow between 1 and 10 Nefm, and just four percent had more than ten Nefm, with only two infinite number of females migrating (between 3: MadeMani and 5: AmaPari, and 15: TocaItup and 16: AragAnto) (upper diagonal of the Table S4).

### Ecological niche modeling for resistance landscape analysis

The performance of the *Podocnemis unifilis* Ecological Niche Model (ENM, Fig S1) showed a high AUC value (0.888). The variables of greatest contribution (percent contribution) for the species’ ENM (i.e., the variables that most explain the geographic distribution of the species) were the bioclimatic variable of the Annual Precipitation average [mm/year] across sub-catchment (water courses only) and the freshwater variable of the Water Flow Length, followed by the vegetation variable that showed some greater relative contribution (permutation contribution), the Leaf Area Index (Table 2).

**Table 2.** Environmental variables used in the Ecological Niche Modeling of the *Podocnemis unifilis* and estimates of relative contributions of the environmental variables.

| Variables | Percent contribution | Permutation importance |
| --- | --- | --- |
| Annual Precipitation* | 38.9 | 34.9 |
| Water Flow Length | 24.8 | 9.8 |
| Precipitation of Driest Month* | 9.7 | 2.6 |
| Net Primary Productivity | 8.1 | 8.5 |
| Leaf Area Index | 6.1 | 12.4 |
| Temperature Seasonality* | 3.9 | 4.6 |
| Probability of occurrence (0-100%) of R horizon* | 2.4 | 4.1 |
| Precipitation of Coldest Quarter* | 2 | 0.5 |
| Clay content mass fraction* | 1.4 | 1.9 |
| Cation exchange capacity | 1 | 1.2 |
| Precipitation Seasonality* | 0.6 | 0.2 |
| Digital elevation model | 0.6 | 6.8 |
| Precipitation of Warmest Quarter* | 0.5 | 0.7 |

\*Variables for near freshwater specific environments across sub-catchment.

### Isolation by distance and by resistance

The Least Cost Paths (LCPs) predicted routes between different rivers upstream of the rapids through low-resistance environments available in the landscape, such as between localities of the Araguaia river (localities 16, 17, 18 and 19) and Tocantins river (20, 21 and 22) (Fig. 3).

**Fig. 3.**
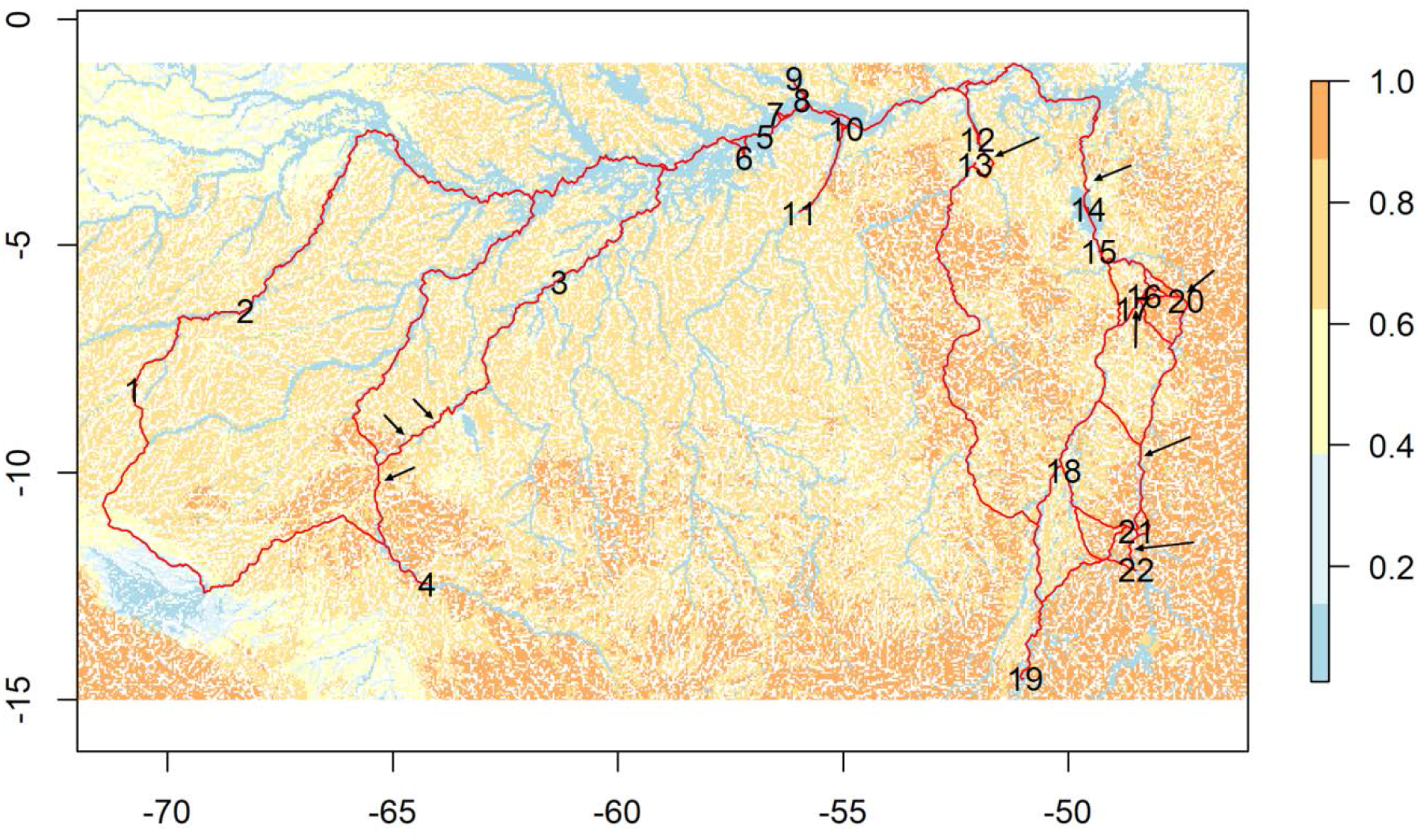
Result from least-cost corridors path (LCP) analyses for the *Podocnemis unifilis* using the Landscape Friction layer considering the Ecological Niche Model for the species with rapids and waterfalls included. Corridors are shown in red and resistance values range from 0 to 1, from lower environmental resistance in blue (corresponding to the freshwater environments with high adequability) to high environmental resistance in orange. Locations of rapids and waterfalls are highlighted with black arrows.

Furthermore, there were routes among the location 4 (MadeCost) crossing the headwaters to the Juruá river and through the Purus river to the Amazon river, and even between Araguaia and Xingu rivers through the upper Xingu where there are no genetic data (Fig. 3).

Although a significant correlation was found between pairwise ΦST and minimum geographic distance (Mantel r = 0.28, p = 0.03) and in-water distance (Mantel r = 0.36, p = 0.02), the greatest correlation was registered by the IBR resistance matrix (r = 0.51, p < 0.001, Table 3).

**Table 3.** Results of the Mantel test, using as response variable the genetic distances (ΦST) with the geographic distance, in water distance, or the resistance distance.

| Predictor | R2 | *p* value |
| --- | --- | --- |
| Geographic | 0.28 | 0.03 |
| Rivers | 0.36 | 0.02 |
| Resistance | 0.51 | < 0.001 |

The multiple regressions MRM showed a similar result of the Mantel within least environmental cost distance (Fig. 4A) with the strongest effect (r^2^ = 0.54, p < 0.0001, Table 4), followed by IBD in-water distance (r^2^ = 0.39, p = 0.004, Fig. 4C, Table 4), and the lowest significant effect by minimum linear distance (r^2^ = 0.31, p = 0.009, Fig. 4B, Table 4). The genetic correlation was high among close localities, but some close localities were less correlated with the more distant ones (Fig. 4D).

**Fig. 4.**
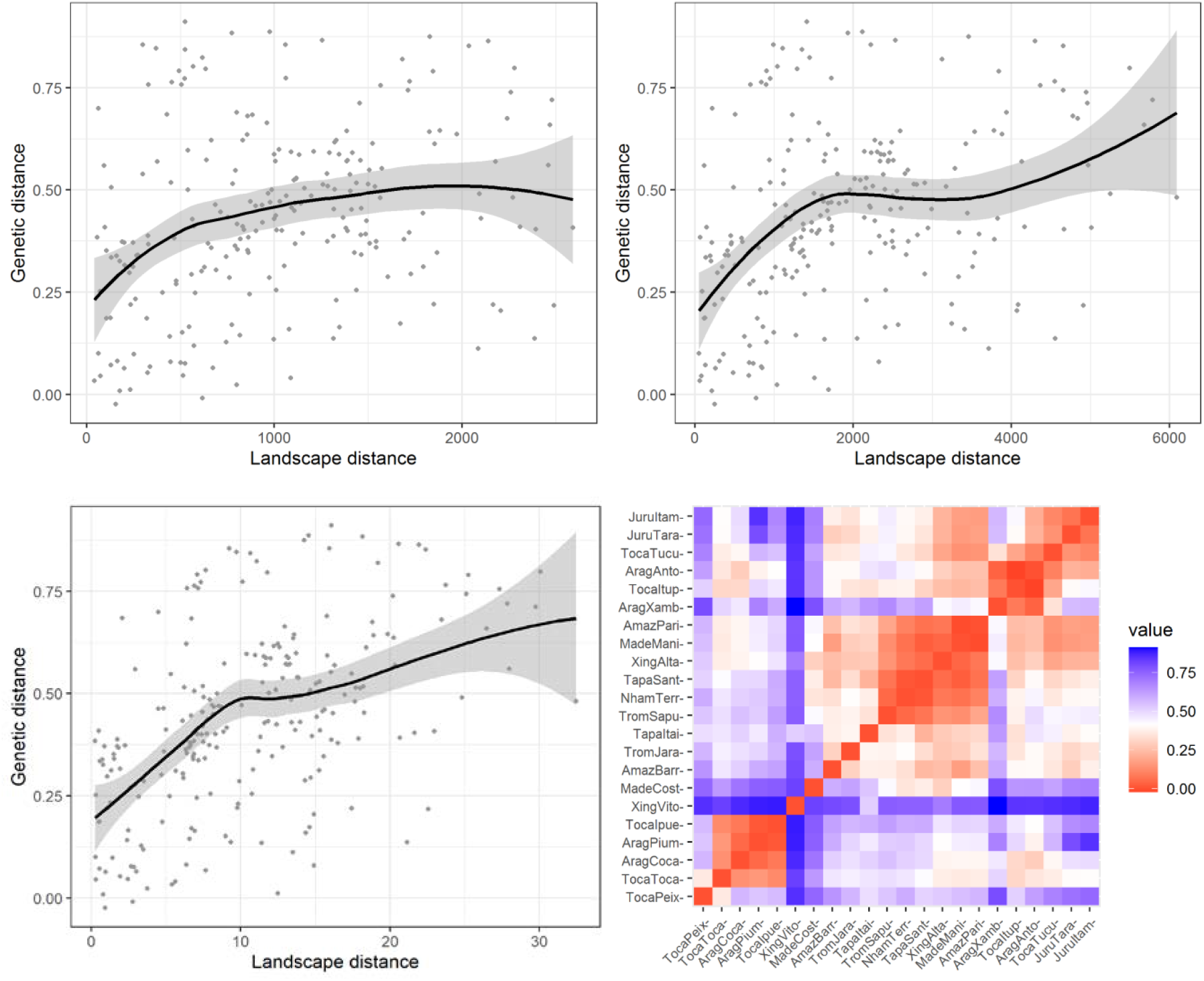
Results from Multiple Matrix Regression with Randomization analyses relating A) resistance, B) geographic, and C) in water to genetic and D) pairwise genetic correlation of the Yellow-spotted River Turtle (*Podocnemis unifilis*).

**Table 4.**
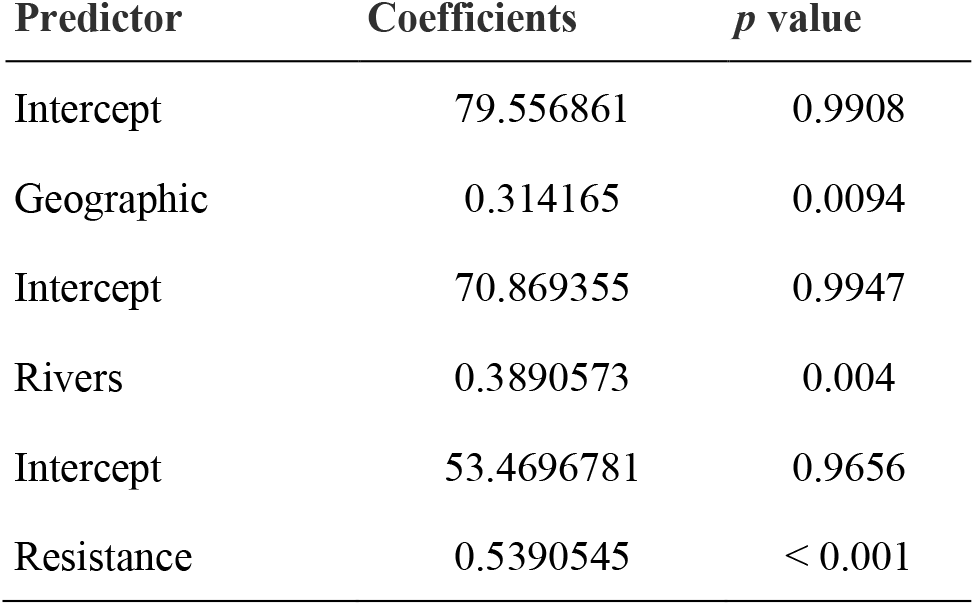
Results of the multiple regressions on dissimilarity matrices (MRM), using as response variable the genetic distances (ΦST) and as explanatory variables the geographic distance, in water distance, or the resistance distance.

| Predictor | Coefficients | *p* value |
| --- | --- | --- |
| Intercept | 79.556861 | 0.9908 |
| Geographic | 0.314165 | 0.0094 |
| Intercept | 70.869355 | 0.9947 |
| Rivers | 0.3890573 | 0.004 |
| Intercept | 53.4696781 | 0.9656 |
| Resistance | 0.5390545 | < 0.001 |

## Discussion

In this study we evaluated genetic structuring of *Podocnemis unifilis* using the mtDNA. We used different landscape factors to explain the genetic variation, testing IBD by linear geographic distance, isolation by in-water distance (i.e., through rivers), and IBR by landscape resistance considering different variables specific to the aquatic environment, including high resistance levels for rapids and waterfalls. We found a heterogeneous genetic diversity for the species, and some genetic structure which seems to follow some watersheds. We found that landscape resistance (IBR) better explains genetic distance than geographic or even following in-water distance (IBD).

### Genetic diversity patterns of Podocnemis unifilis

Analyzing the variability indices, we found different levels of genetic diversity into the localities of *P. unifilis* along the Brazilian Amazonia, from low levels of haplotypic diversity (between 0.000 and 0.500), intermediary (between 0.694 and 0.749) to high levels (0.814 to 1.000), with an overall average of 0.712. These results agree with the average values findings for other species of the genus also within the mtDNA control region (0.51 for *P. erythrocephala;* Santos et al. 2016; Michels and Vargas-Ramirez 2018, 0.65 for *P. expansa;* Pearse et al. 2006, and 0.77 for *P. sextuberculata;* Viana et al. 2017). When we analyzed these indices by biological groups, we found higher values, between 0.776 (for the BG4, from Araguaia-Tocantins drainage) and 0.886 (for the BG3, basically from the Xingu river upper the waterfalls), with an average of 0.821, showing that within the biological group’s individuals present greater diversity in relation to this comparison between localities. This may indicate a general pattern of a high level of diversity species, with some localities as an exception.

The low levels of haplotypic diversity found in five localities (0.000 in 18: AragPium; 0.318 in 2: JuruItam; 0.433 in 4: MadeCost, 0.439 in 1: JuruTara, and 0.500 in 6: AmazBarr) need to be carefully noticed, considering that the genetic diversity is directly related to the size and the fitness of the population. Furthermore, the levels of genetic diversity are one of the three necessary indicators for the conservation of biodiversity by the IUCN that are also related with the area of occupancy, extent of occurrence, number of mature individuals, generation length, reduction in the number of mature etc. (IUCN 2001; Frankham 2012; Frankham et al. 2014).

### Population structure and gene flow

The analysis of molecular variance showed a high degree of subdivision between the localities verified in most pairwise comparisons (Table S4). The haplotype network showed several (>20% of the sampled individuals) unique haplotypes with different of branch lengths caused by the migration of individuals between localities (Salzburger et al. 2011). On the other hand, there were many shared haplotypes (up to 61 individuals in the same haplotype), indicating some ancestral haplotypes, and possibly recent expansion, specially at localities within Araguaia-Tocantins interfluve as highlighted by the neutrality tests (six of the nine sites with significant values of *D* [from −1.934 to −0.774] and/or *Fs* [from - 834.057 to 101.973]; Table 1).

The Araguaia-Tocantins river basin was formed in the Pleistocene around 240,000 years ago (Valente et al. 2013), has relatively stationary geographic features presenting more permanent physical obstacles and is in an ecotonal zone between the Amazonia and the Cerrado (IBGE 2019). Such characteristics made this basin an excellent natural setting to test biogeographic hypotheses from ancient and/or recent events such as recent quaternary cycles and even uplift of plateaus or marine transgressions (Rocha et al. 2015; Guarnizo et al. 2016).

Studies with aquatic organisms have correlated the more recently formation of the Araguaia-Tocantins basin (Regarding the formation of the Amazon river basin established during the Miocene) and the fact that it does not drain directly into the Amazon river basin (Goulding et al. 2003; Farias et al. 2019; Hoorn et al. 2022) as a cause of speciation (e.g., dolphins; Hrbek et al. 2014), or of population genetic differences due to geographic distance (e.g., arapaima; Farias et al. 2019), and perhaps explain this possible recent expansion found in our data.

### Isolation by distance and by resistance

Within BG1 and BG2there are genetically very similar individuals from different localities of the Solimões-Amazon system, mainly in the lowland Amazonian region. This pattern has also been found for other aquatic animals, such as fishes (Hrbek et al. 2005; Willis et al. 2012b; Ochoa et al. 2015), stingrays (Frederico et al. 2012), and even turtles (Pearse et al. 2006) to which the main populations of the Amazon river valley (white water floodplain system; Junk et al. 2012) did not show genetic structure among them. The Giant Amazon River Turtle (*P. expansa*), in general, showed a pattern of structuring between sub-basins (genetic structure among drainages of different rivers and watersheds) with significant isolation by distance (Pearse et al. 2006), in agreement with *P. unifilis* found by Escalona et al. (2009). Although we have also found genetic structure for *P. unifilis* between different sub-basins, the geographic distance or even the distance in-water did not represent the best proxy for genetic isolation in this species considering the mitochondrial DNA marker investigated here.

Considering that the variables that most explain the geographic distribution of *P. unifilis* were the average annual rainfall and the length of the water flow, the influence of landscape resistance on the movement of individuals may be related to reproductive adaptation, or at least influence the start of the nesting period, which depends on the decrease in water level and varies according to the regional hydrological cycle(Ferrara et al. 2017).

Another variable that presented a greater relative contribution, but through a slightly negative relationship, was the Leaf Area Index (LAI). This variable expresses the rate of vegetation growth correlated with productivity, that is, the gas exchange from the leaf to the canopy (Breda 2003). LAI also reflect on reproductive aspects of the Yellow-Spotted Amazon River Turtle, since despite being able to adapt its nesting behaviour in different environments and substrates (beaches, floodplains, ravines, sand, clay; Andrade, 2012, 2015), especially in areas with undergrowth, or close to forests in overgrown places, it is more common to see nests in beaches (places with low vegetation productivity), where the adult individuals are more vulnerable and is easier to hunting (Erickson et al. 2020; Balestra and Lacava 2020; Fagundes et al. 2021).As this species has greater terrestrial mobility than other species (Vogt 2008; Andrade, 2012; Ferrara et al. 2017), was found evidence that this species can bypass obstacles, possibly connecting populations from different rivers when the landscape is suitable for its occurrence (Escalona et al. 2009). In agreement, we found that the resistance of the landscape, considering specific variables of the freshwater environment, better explains the genetic differentiation of this species. Furthermore, the rapids and waterfalls (including their magnitude levels) tested here function as barriers with different levels of permeability for this species, with low permeability between locations with greater genetic structuring such as the Guaporé, Xingu and Tocantins rivers.

### Implications for Conservation

Except for some locations in the Araguaia-Tocantins drainage, most localities sampled are evolving under the hypothesis of selective neutrality with mtDNA (Hartl and Clark 1989), although analysis for the same species using microsatellites data showed signs of reduction in population size in a more recent past in Western Amazonia (Escalona et al. 2009). This potentially indicates that *P. unifilis* individuals can increase genetic diversity if the reduction in population numbers does not persist (Frankham et al. 2008). But, considering that the main threat is overexploitation by the illegal trade for consumption, associated with high habitat reduction and degradation, current conservation efforts may be insufficient (Ferrara et al. 2017; Chaves et al. 2021), at least to guarantee the survival of this species in some localities. As an example, community-based turtle conservation projects (such as the Pé-de-pincha Program; Andrade 2022) that range many localities and different river channels in the Amazon, providing stable or increasing populations of *P. unifilis* in different areas and watersheds (Andrade et al. 2022). Furthermore, for successful management and conservation for organisms that have a long-life cycle, such as chelonians, it is also necessary to protect all life stages of the species (Congdon, Dunham and van Loben Sels 1993; Frankham et al. 2008; Eisemberg et al. 2019) and all areas of occurrence, mainly because the main conservation efforts focus on reproductive areas and often neglect feeding areas, as important as for the life history of these organisms.

Since the main locations observed with the highest degree of differentiation are above waterfalls in the Guaporé, Xingu and Tocantins rivers (4: MadeCost, 13: XingVito, 14: TocaTucu, 20: TocaToca and 22: TocaPeix), the information contained in this article can be taken into consideration for species conservation. Currently, several hydroelectric plants are under construction/operating or planned in the region (Winemiller et al. 2016; Fagundes et al. 2021). Those dams rupture the connectivity of rivers longitudinally, and the main channels with riparian zones and floodplains making the migration of freshwater turtles more difficult, reducing its adaptive capacity (Fagundes et al. 2018). Furthermore, it changes the hydrological regime flooding watercourses, or at least decreasing the exposure time of the turtles’ nesting sandbanks, impacting incubation rate and reproductive success (Eisemberg et al. 2016; Ferrara et al. 2017). In this context, our results can help in decisions about dams, improving the knowledge of such impacts, since the many current Brazilian Environmental Impact Studies (EIA - Estudo de Impacto Ambiental) tend to minimize or ignore significant impacts (Fearnside 2015) through weak technical and/or scientific support (Ritter et al. 2017).

For instance, we found that the most localities have at least one migrant between leastwise one other locality (Table S4) using an estimated metric to analyze the number of female migrants per generation (Nefm), indicating that genetic divergence is being contained and those populations presented restricted geneflow (Hartl and Clark 1989). A single exception was the locality above the ‘Volta Grande’ waterfalls of Xingu river (13: XingVito), also showed by BAPS results, with less than one female migrating per generation compared to all other localities. There is no estimate for the generation time for *P. unifilis*, but for the sister clade *P. lewyana* (Vargas-Ramírez et al. 2008) is ten years (Páez et al. 2015). Thus, we can predict that a female individual of the yellow-spotted female river turtle would take at least 46 years (0.22 Nefm) to cross the waterfalls of the Xingu river (between localities 13: XingVito and 12: XingAlta). Considering that would require a large amount of energy and the turtle would spend a large part of its life to cross the ‘Volta Grande’ by the river, we can characterize these waterfalls as a great barrier for the dispersal of *P. unifilis*.

This region of the Xingu ‘Volta Grande’ is composed of huge morphological complexes rapids, comprising a unique environment that plays an important role in Amazonian aquatic endemism and biodiversity (Sawakuchi et al. 2015). Analyzing the least cost paths (LCPs), the populations from the Xingu river possibly used a corridor to the Araguaia river (Fig. 3). Therefore, we suggest that possibly non-sampled localities of the upper Xingu river may be more related to Araguaia-Tocantins samples due to the environmental resistance barrier in the middle Xingu revealed by the ENM.

## Conclusions

This was the first study to assess landscape genetics for a widely distributed neotropical turtle spanning different river basins. We found heterogeneous levels of genetic diversity with caveat for species conservation. *Podocnemis unifilis* presents complex geographic patterns, with localities grouped biologically, and localities significantly different from each other. We verified that the resistance of the landscape influences the displacement of individuals by aquatic, vegetational, biological and geomorphological variables. Efforts to conserve the species need to be applied throughout its distribution considering landscape genetics, not just in Brazil.

## Author contributions

MAPA designed the study with AFM, CDR and IPF. WPOJ and RCV were mainly responsible for field work. PCMA collected samples in the middle and lower Amazon river and in the Juruá, Madeira, Andirá, Nhamundá and Trombetas rivers. MNSV and MAPA obtained most DNA sequences. MAPA, AFM, CDR and MNSV generated and MAPA, AFM, CDR and IPF analyzed the data. MAPA wrote the manuscript with contributions of all.

## Data availability

GenBank accession sequences ID

## Acknowledgments

The fieldwork authorization was granted by the Instituto Brasileiro do Meio Ambiente e dos Recursos Naturais Renováveis (IBAMA [Brazilian Institute of the Environment and Renewable Natural Resources]) and the Instituto Chico Mendes de Conservação da Biodiversidade (ICMBio [Chico Mendes Institute of Conservation of Biodiversity]) (licenses number: 12388-4, 49695-1 and 49289). For the fieldwork groups of Laboratório de Biotecnologia and Quelônios e Crocodilianos da Região Norte from the Universidade Federal do Tocantins (UFT [Federal University of Tocantins]), and Pé-de-Pincha Project from the Universidade Federal do Amazonas (UFAM [Federal University of Amazonas]).

## Supplementary Information

**Table S1.** Occurrence records of *Podocnemis unifilis* used to generate Ecological Niche Modeling.

(Indexed)

**Table S2.** Rapids and water-falls level values included in the resistance landscape scaled for 0.5 to 0.85 proportional to their extension size. Extension sizes are in meters.

| ID | Rapids and Waterfalls | River | Longitude | Latitude | Extension | Resistance |
| --- | --- | --- | --- | --- | --- | --- |
| A | Misericórdia | Madeira | -65.2918 | -10.2282 | 5,000 | 0.51 |
| B | Caldeirão | Madeira | -64.63793 | -9.25317 | 4,070 | 0.51 |
| C | Teotônio | Madeira | -64.06406 | -8.85804 | 2,700 | 0.50 |
| D | Santo Antônio | Madeira | -63.93845 | -8.79293 | 2,000 | 0.50 |
| E | Volta Grande do Xingu | Xingu | -51.71046 | -3.3959 | 130,000 | 0.85 |
| F | Taboca | Tocantins | -49.6479 | -3.83211 | 11,000 | 0.52 |
| G | Santa Isabel | Araguaia | -48.39872 | -6.14813 | 14,000 | 0.53 |
| H | São João | Araguaia | -48.42459 | -6.22729 | 2,350 | 0.50 |
| I | Remanso dos Botos | Araguaia | -48.37685 | -6.35349 | 5,000 | 0.51 |
| J | São Miguel | Araguaia | -48.50944 | -6.35538 | 6,156 | 0.51 |
| K | Santa Rita | Tocantins | -47.47321 | -5.7673 | 3,000 | 0.50 |
| L | Lageado | Tocantins | -48.37305 | -9.75562 | 5,400 | 0.51 |
| M | Arquipélago do Tropeço | Tocantins | -48.50666 | -12.0929 | 6,800 | 0.51 |

**Table S3.**
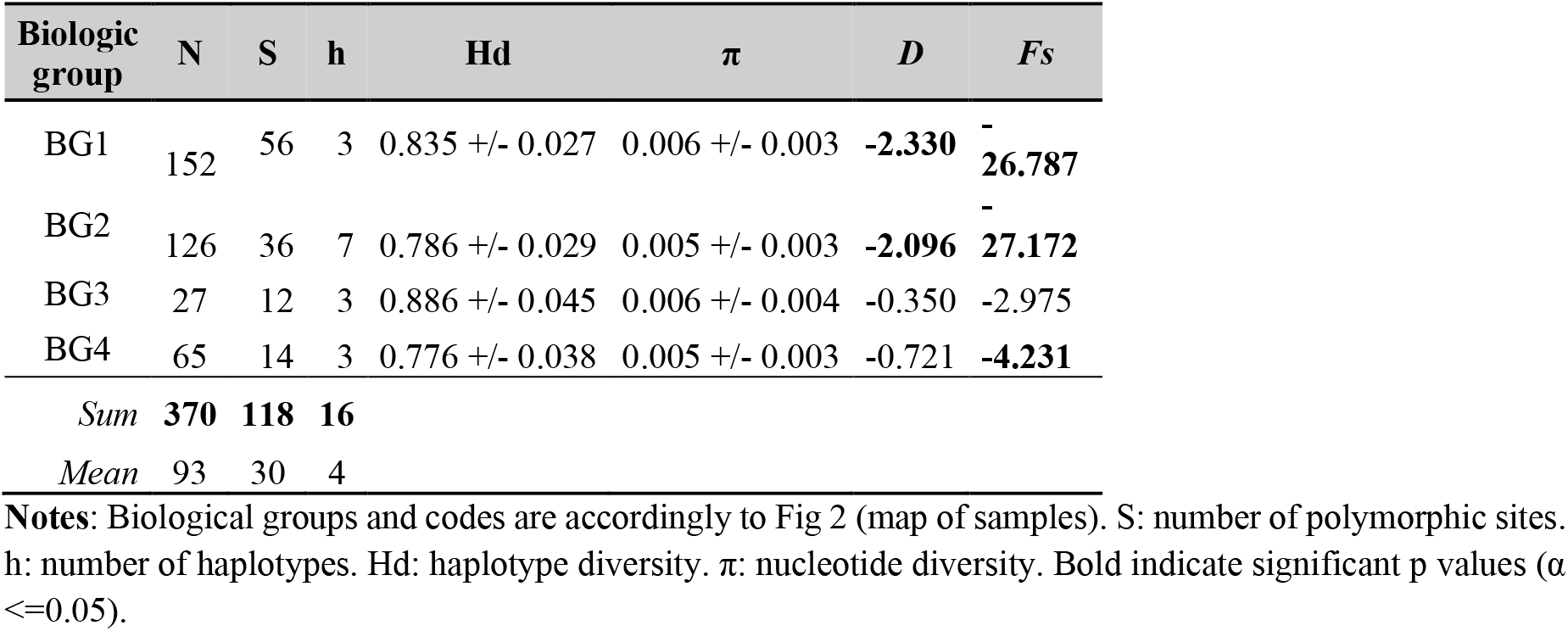
Genetic diversity indices of the *Podocnemis unifilis* biological groups from BAPS.

**Table S4.**
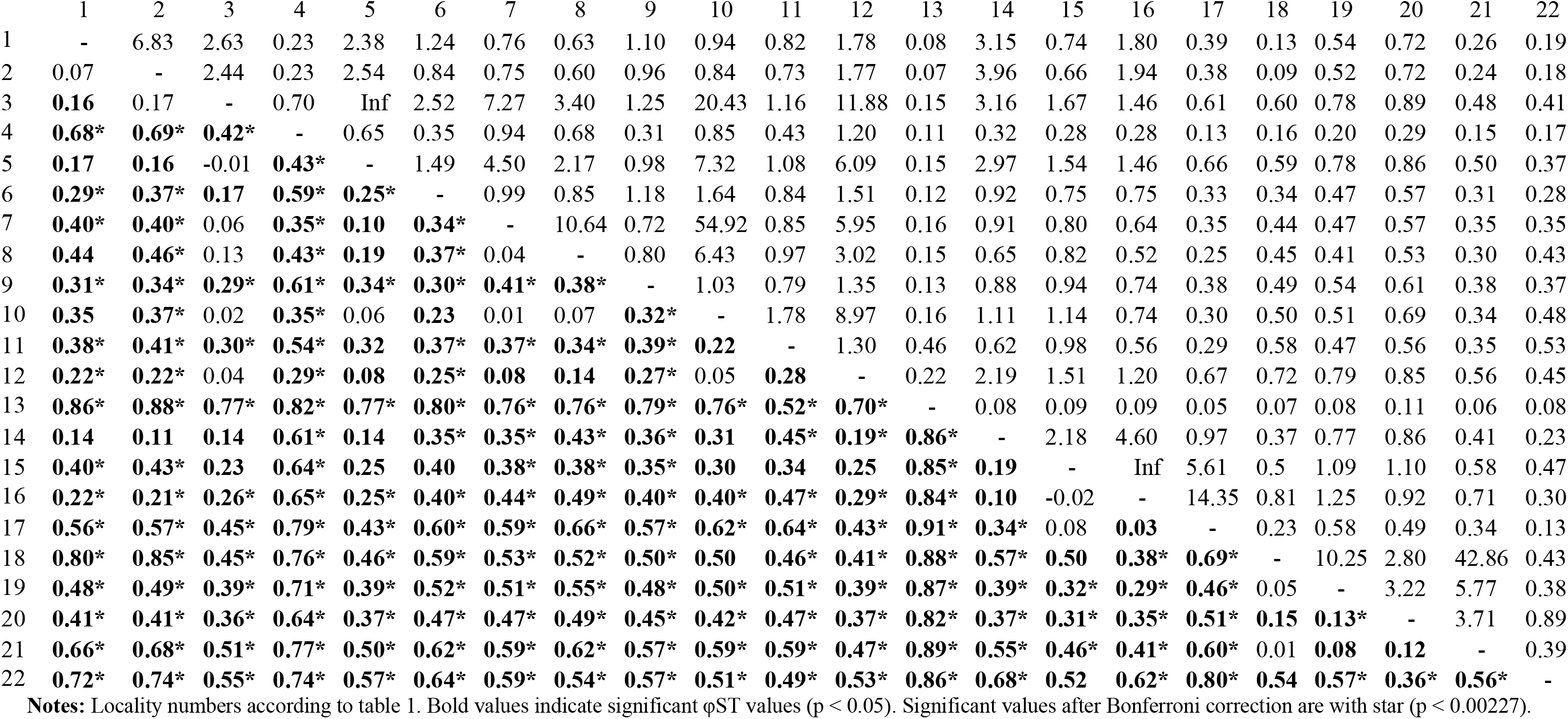
Pairwise φST (lower diagonal) and number of migrating *Podocnemis unifilis* females per generation between localities studied (upper diagonal).

|  | 1 | 2 | 3 | 4 | 5 | 6 | 7 | 8 | 9 | 10 | 11 | 12 | 13 | 14 | 15 | 16 | 17 | 18 | 19 | 20 | 21 | 22 |
| --- | --- | --- | --- | --- | --- | --- | --- | --- | --- | --- | --- | --- | --- | --- | --- | --- | --- | --- | --- | --- | --- | --- |
| 1 | - | 6.83 | 2.63 | 0.23 | 2.38 | 1.24 | 0.76 | 0.63 | 1.10 | 0.94 | 0.82 | 1.78 | 0.08 | 3.15 | 0.74 | 1.80 | 0.39 | 0.13 | 0.54 | 0.72 | 0.26 | 0.19 |
| 2 | 0.07 | - | 2.44 | 0.23 | 2.54 | 0.84 | 0.75 | 0.60 | 0.96 | 0.84 | 0.73 | 1.77 | 0.07 | 3.96 | 0.66 | 1.94 | 0.38 | 0.09 | 0.52 | 0.72 | 0.24 | 0.18 |
| 3 | **0.16** | 0.17 | - | 0.70 | Inf | 2.52 | 7.27 | 3.40 | 1.25 | 20.43 | 1.16 | 11.88 | 0.15 | 3.16 | 1.67 | 1.46 | 0.61 | 0.60 | 0.78 | 0.89 | 0.48 | 0.41 |
| 4 | **0.68\*** | **0.69\*** | **0.42\*** | - | 0.65 | 0.35 | 0.94 | 0.68 | 0.31 | 0.85 | 0.43 | 1.20 | 0.11 | 0.32 | 0.28 | 0.28 | 0.13 | 0.16 | 0.20 | 0.29 | 0.15 | 0.17 |
| 5 | **0.17** | **0.16** | -0.01 | **0.43\*** | - | 1.49 | 4.50 | 2.17 | 0.98 | 7.32 | 1.08 | 6.09 | 0.15 | 2.97 | 1.54 | 1.46 | 0.66 | 0.59 | 0.78 | 0.86 | 0.50 | 0.37 |
| 6 | **0.29\*** | **0.37\*** | **0.17** | **0.59\*** | **0.25\*** | - | 0.99 | 0.85 | 1.18 | 1.64 | 0.84 | 1.51 | 0.12 | 0.92 | 0.75 | 0.75 | 0.33 | 0.34 | 0.47 | 0.57 | 0.31 | 0.28 |
| 7 | **0.40\*** | **0.40\*** | 0.06 | **0.35\*** | **0.10** | **0.34\*** | - | 10.64 | 0.72 | 54.92 | 0.85 | 5.95 | 0.16 | 0.91 | 0.80 | 0.64 | 0.35 | 0.44 | 0.47 | 0.57 | 0.35 | 0.35 |
| 8 | **0.44** | **0.46\*** | 0.13 | **0.43\*** | **0.19** | **0.37\*** | 0.04 | - | 0.80 | 6.43 | 0.97 | 3.02 | 0.15 | 0.65 | 0.82 | 0.52 | 0.25 | 0.45 | 0.41 | 0.53 | 0.30 | 0.43 |
| 9 | **0.31\*** | **0.34\*** | **0.29\*** | **0.61\*** | **0.34\*** | **0.30\*** | **0.41\*** | **0.38\*** | - | 1.03 | 0.79 | 1.35 | 0.13 | 0.88 | 0.94 | 0.74 | 0.38 | 0.49 | 0.54 | 0.61 | 0.38 | 0.37 |
| 10 | **0.35** | **0.37\*** | 0.02 | **0.35\*** | 0.06 | **0.23** | 0.01 | 0.07 | **0.32\*** | - | 1.78 | 8.97 | 0.16 | 1.11 | 1.14 | 0.74 | 0.30 | 0.50 | 0.51 | 0.69 | 0.34 | 0.48 |
| 11 | **0.38\*** | **0.41\*** | **0.30\*** | **0.54\*** | **0.32** | **0.37\*** | **0.37\*** | **0.34\*** | **0.39\*** | **0.22** | - | 1.30 | 0.46 | 0.62 | 0.98 | 0.56 | 0.29 | 0.58 | 0.47 | 0.56 | 0.35 | 0.53 |
| 12 | **0.22\*** | **0.22\*** | 0.04 | **0.29\*** | **0.08** | **0.25\*** | **0.08** | **0.14** | **0.27\*** | 0.05 | **0.28** | - | 0.22 | 2.19 | 1.51 | 1.20 | 0.67 | 0.72 | 0.79 | 0.85 | 0.56 | 0.45 |
| 13 | **0.86\*** | **0.88\*** | **0.77\*** | **0.82\*** | **0.77\*** | **0.80\*** | **0.76\*** | **0.76\*** | **0.79\*** | **0.76\*** | **0.52\*** | **0.70\*** | - | 0.08 | 0.09 | 0.09 | 0.05 | 0.07 | 0.08 | 0.11 | 0.06 | 0.08 |
| 14 | **0.14** | **0.11** | **0.14** | **0.61\*** | **0.14** | **0.35\*** | **0.35\*** | **0.43\*** | **0.36\*** | **0.31** | **0.45\*** | **0.19\*** | **0.86\*** | - | 2.18 | 4.60 | 0.97 | 0.37 | 0.77 | 0.86 | 0.41 | 0.23 |
| 15 | **0.40\*** | **0.43\*** | **0.23** | **0.64\*** | **0.25** | **0.40** | **0.38\*** | **0.38\*** | **0.35\*** | **0.30** | **0.34** | **0.25** | **0.85\*** | **0.19** | - | Inf | 5.61 | 0.5 | 1.09 | 1.10 | 0.58 | 0.47 |
| 16 | **0.22\*** | **0.21\*** | **0.26\*** | **0.65\*** | **0.25\*** | **0.40\*** | **0.44\*** | **0.49\*** | **0.40\*** | **0.40\*** | **0.47\*** | **0.29\*** | **0.84\*** | **0.10** | -0.02 | - | 14.35 | 0.81 | 1.25 | 0.92 | 0.71 | 0.30 |
| 17 | **0.56\*** | **0.57\*** | **0.45\*** | **0.79\*** | **0.43\*** | **0.60\*** | **0.59\*** | **0.66\*** | **0.57\*** | **0.62\*** | **0.64\*** | **0.43\*** | **0.91\*** | **0.34\*** | 0.08 | **0.03** | - | 0.23 | 0.58 | 0.49 | 0.34 | 0.13 |
| 18 | **0.80\*** | **0.85\*** | **0.45\*** | **0.76\*** | **0.46\*** | **0.59\*** | **0.53\*** | **0.52\*** | **0.50\*** | **0.50** | **0.46\*** | **0.41\*** | **0.88\*** | **0.57\*** | **0.50** | **0.38\*** | **0.69\*** | - | 10.25 | 2.80 | 42.86 | 0.43 |
| 19 | **0.48\*** | **0.49\*** | **0.39\*** | **0.71\*** | **0.39\*** | **0.52\*** | **0.51\*** | **0.55\*** | **0.48\*** | **0.50\*** | **0.51\*** | **0.39\*** | **0.87\*** | **0.39\*** | **0.32\*** | **0.29\*** | **0.46\*** | 0.05 | - | 3.22 | 5.77 | 0.38 |
| 20 | **0.41\*** | **0.41\*** | **0.36\*** | **0.64\*** | **0.37\*** | **0.47\*** | **0.47\*** | **0.49\*** | **0.45\*** | **0.42\*** | **0.47\*** | **0.37\*** | **0.82\*** | **0.37\*** | **0.31\*** | **0.35\*** | **0.51\*** | **0.15** | **0.13\*** | - | 3.71 | 0.89 |
| 21 | **0.66\*** | **0.68\*** | **0.51\*** | **0.77\*** | **0.50\*** | **0.62\*** | **0.59\*** | **0.62\*** | **0.57\*** | **0.59\*** | **0.59\*** | **0.47\*** | **0.89\*** | **0.55\*** | **0.46\*** | **0.41\*** | **0.60\*** | 0.01 | **0.08** | **0.12** | - | 0.39 |
| 22 | **0.72\*** | **0.74\*** | **0.55\*** | **0.74\*** | **0.57\*** | **0.64\*** | **0.59\*** | **0.54\*** | **0.57\*** | **0.51\*** | **0.49\*** | **0.53\*** | **0.86\*** | **0.68\*** | **0.52** | **0.62\*** | **0.80\*** | **0.54** | **0.57\*** | **0.36\*** | **0.56\*** | - |

**Notes:** Locality numbers according to table 1. Bold values indicate significant ϕST values ( $p < 0.05$ ). Significant values after Bonferroni correction are with star ( $p < 0.00227$ ).

**Figure S1.**
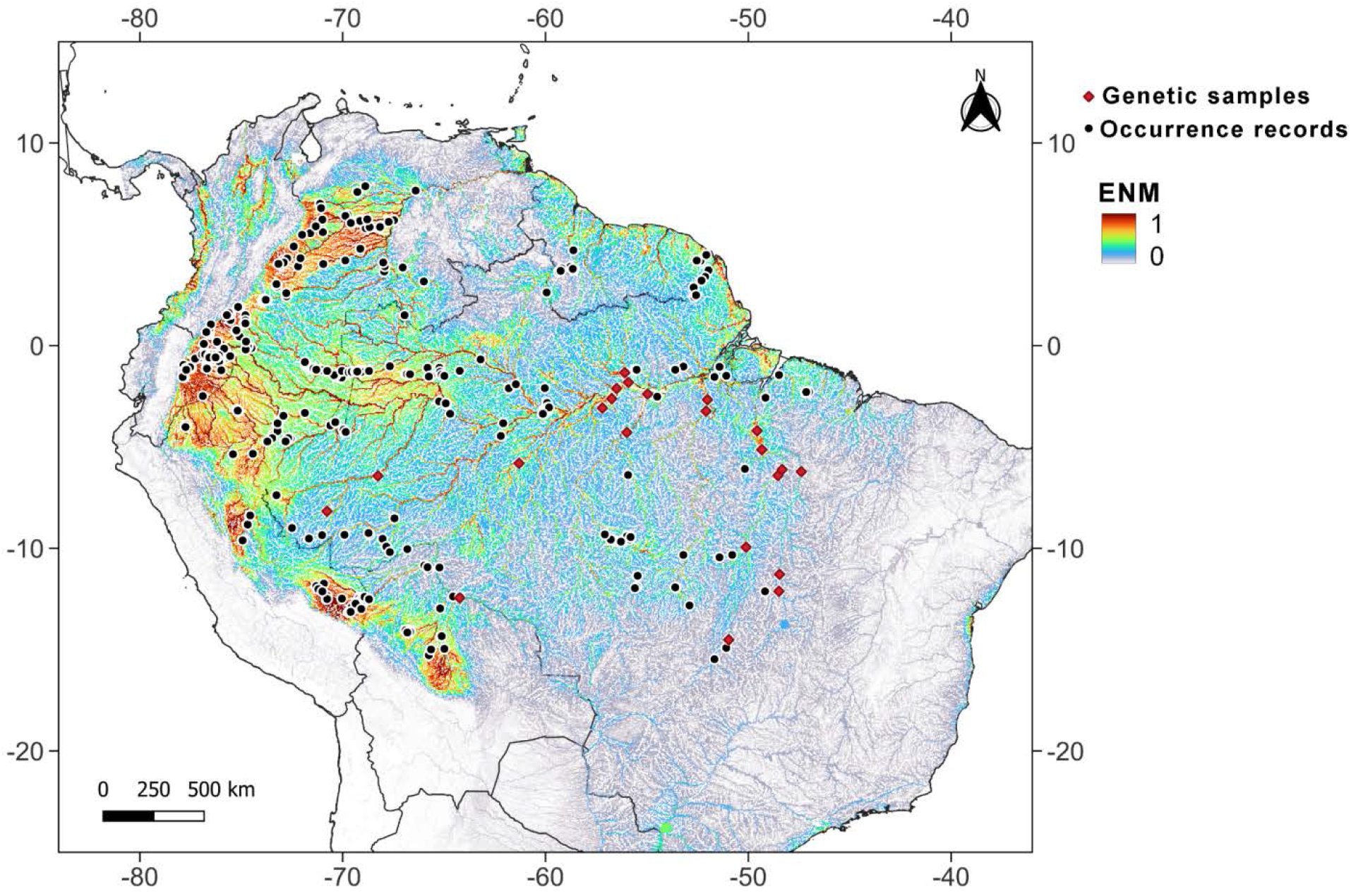
Ecological niche modeling (ENM) for *Podocnemis unifilis* ranging from 0 in cold colors (for low environmental suitability for species occurrence) to 1 in hot colors (for high environmental suitability for species occurrence). Black dots are the occurrence records used to generate the ENM without genetic data and red diamonds are the occurrence records used to generate the ENM with genetic sampling

**Figure S2.**
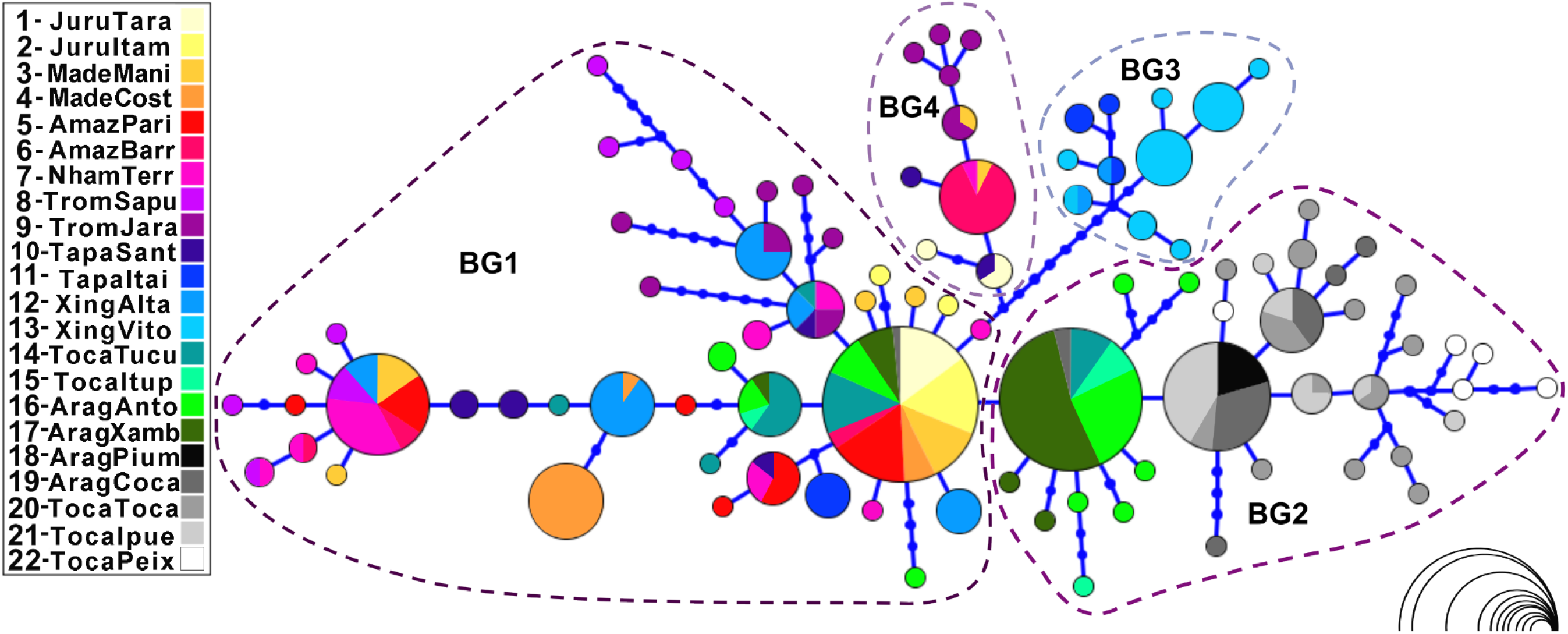
Haplotype network of *Podocnemis unifilis* based on the mtDNA control region. The circle size represents the number of individuals sampled (ranging from 1 to 61 according to the scale at the bottom right).

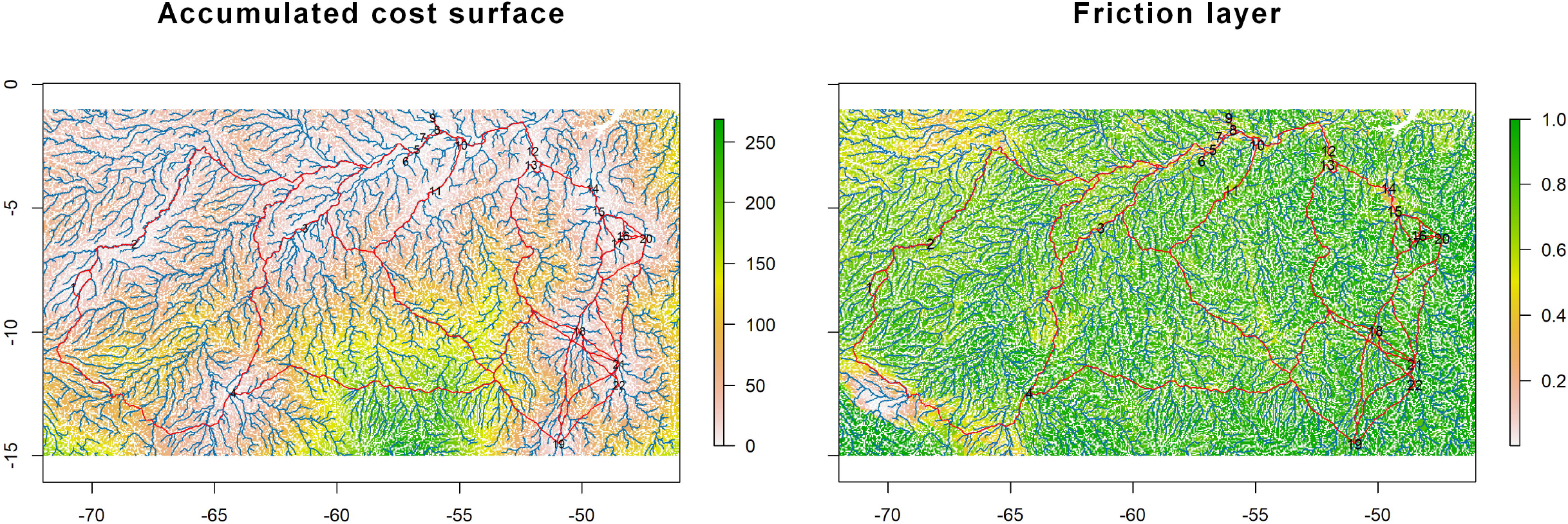

